# Structural constraints to information flow within cortical circuits: a TMS/EEG-dMRI study

**DOI:** 10.1101/026419

**Authors:** Enrico Amico, Olivier Bodart, Olivia Gosseries, Lizette Heine, Mario Rosanova, Pieter Van Mierlo, Charlotte Martial, Marcello Massimini, Daniele Marinazzo, Steven Laureys

## Abstract

Transcranial magnetic stimulation (TMS) has been used for more than 20 years to investigate brain function by perturbing and observing the consequent behavioral, pathophysiological and electrophysiological modulations. These latter, mainly measured by high-density electroencephalography (hd-EEG), revealed signatures of the functional organization in a brain network. In order to unveil the nature and the underlying mechanism of these signatures, we here mapped TMS-induced hd-EEG changes onto changes in information flow and brain structural architecture, using multimodal modeling of source reconstructed TMS/hd-EEG recordings and diffusion magnetic resonance imaging (dMRI) tractography in a cohort of awake healthy volunteers. We observed that the relationship between information flow and structural connections depend on the stimulation site and on the frequency of the TMS-induced brain rhythms. These findings highlight the importance of taking into account the dynamics of different local oscillations when investigating the mechanisms for integration and segregation of information in the human brain. Our whole-brain analysis sheds light on the function-structure organization of the brain network after TMS, and on the huge variety of information contained in it. TMS/EEG dMRI directed functional connectivity structural connectivity structure-function brain information flow

## Introduction

Transcranial magnetic stimulation (TMS) has been used for more than 20 years to investigate connectivity and plasticity in the human cortex. By combining TMS with high-density electroencephalography (hd-EEG), one can stimulate any cortical area and measure the effects produced by this perturbation in the rest of the cerebral cortex (Ilmoniemi et al., 1997; Komssi and Kähkönen, 2006). It has been shown that cortical potentials elicited by TMS stimulation (TMS-evoked potentials, i.e. TEPs) last for up to 600 ms in normal wakefulness, during their spread from the area of stimulation to remote interconnected brain areas (Bonato et al., 2006; Lioumis et al., 2009). To date, TMS/EEG recordings have provided new insights on the whole brain cortical excitability with reasonable spatial and excellent temporal resolution (Rogasch and Fitzgerald, 2013).

The amount of information contained in the hd-EEG response to TMS has appeared to contain inner signatures of the functional organization in a brain network. A recent study (Rosanova et al., 2009) in healthy awake subjects showed that TMS can also induce EEG oscillations at different frequencies. The TMS pulse gives rise to different connected cortical regions in the brain, generating a complex EEG pattern composed of strong fluctuations at the “resonant”(natural) frequency of the stimulated area. These oscillations are thought to reflect neurophysiological activity that is transiently elicited by the TMS pulse and possibly engaged through brain connections (Rosanova et al., 2009).

The study of TEPs has increased our understanding of cortical processing both in health (Massimini et al., 2005; Ferrarelli et al., 2010) and disease (Rosanova et al., 2012; Ragazzoni et al., 2013; George et al., 2000; Gosseries et al., 2015). For instance, from compressing the information given by TMS, Casali et al. defined an empirical measure of brain complexity, i.e. the perturbational complexity index (PCI). They demonstrated that this index can reliably discriminate between different physiological, pharmacological, and pathological levels of consciousness (Casali et al., 2013).

Recently, researchers have started to investigate how the TMS/hd-EEG perturbation might be constrained and shaped by brain structure, either by exploring the correlation between TMS-induced interhemispheric signal propagation and neuroanatomy (Groppa et al., 2013; Voineskos et al., 2010), or by improving the modeling of the TMS-induced electric field using realistic neural geometry (De Geeter et al., 2015; Bortoletto et al., 2015). Besides, it has lately been shown that cortical networks derived from source EEG connectivity partially reflects both direct and indirect underlying white matter connectivity in a broad range of frequencies (Chu et al., 2014).

In this respect, the development of diffusion magnetic resonance imaging (dMRI) (Basser, 1995) might add information on the structural architecture of the brain (Catani et al., 2002). The application of deterministic and probabilistic tractography methods allows for the spatial topography of the white matter, which represents bundles coherently organized and myelinated axons (Song et al., 2002). The output of tractography algorithms permits anatomically plausible visualization of white matter pathways (Hofer and Frahm, 2006) and has led to reliable quantification (Voineskos et al., 2009) of structural connections between brain regions (i.e. the brain connectome (Sporns et al., 2005; Bullmore and Sporns, 2009)).

The purpose of this paper is to investigate EEG changes of information flow in the brain induced by TMS from both a functional and structural perspective, using multimodal modeling of source reconstructed TMS/hd-EEG recordings and dMRI tractography. The study of information transfer after the perturbation can possibly help in understanding the structure-function modulation caused by TMS (i.e. the extent to which TMS-induced EEG dynamics is constrained by white matter pathways) and the specific frequency bands of the involved brain regions. Functional and structural connectivity in the brain are known to be closely correlated (Bullmore and Sporns, 2009; Honey et al., 2009; Chu et al., 2014), but their interactions remain only poorly understood (Honey et al., 2010).

Taking the aforementioned recent findings as a starting point, we here aim to assess: 1) if the extent to which information transfer changes in a cortical region, as a consequence of the induced perturbation, is related to the number of fiber pathways passing through it (Chu et al., 2014); 2) whether the temporal variability of the response to TMS has specific spectral EEG signatures (Rosanova et al., 2009); 3) the role of these “natural frequencies” in the flow spread during TMS and in the structure-function interactions (Massimini et al., 2005; Casali et al., 2013; Rosanova et al., 2009).

We will first present the processing pipelines for TMS-EEG and dMRI data. Second, the mathematical methodology for the evaluation of the information flow between brain regions and its correlation with the structural connectome will be presented. Finally, results obtained in a cohort of healthy volunteers (*n* = 14) will be presented and discussed.

## Materials and Methods

### TMS/hd-EEG recordings

#### Acquisition and preprocessing

TMS/hd-EEG data were acquired in 14 healthy awake adults (6 males and 8 females, age range 23-37 years) as published elsewhere (Casali et al., 2013; Rosanova et al., 2012). In brief, subjects were lying with eyes open looking at a fixation point on a screen. All participants gave written informed consent and underwent clinical examinations to rule out any potential adverse effect of TMS. The TMS/hd-EEG experimental procedure, approved by the Local Ethical Committee of University of Liège, was performed using a figure-of-eight coil driven by a mobile unit (eXimia TMS Stimulator, Nexstim Ltd., Finland), targeting two cortical areas (left precuneus and left premotor) for at least 200 trials. These areas were selected for the following reasons: (i) they are easily accessible and far from major head or facial muscles whose activation may affect EEG recordings and (ii) previous TMS/EEG studies have been successfully performed in these areas during wakefulness (Rosanova et al., 2009).

The left precuneus and left premotor targets were identified on the subjects 3D T1 brain scan and reached through the neuronavigation system (NBS, Nexstim Ltd, Finland) using stereoscopic infrared tracking camera and reflective sensors on the subject’s head and the stimulating coil. A stimulation target was chosen in the middle of the area, and we modified slightly its position, as well as the stimulation parameters (intensity, angle, direction) in order to avoid artifacts and get the best response (i.e. the higher signal to noise ratio). Stability of the coil position was assured by using an aiming device allowing the stimulation only when the deviation from the target was less than 2 mm. The intensity was chosen in order to assure an induced electrical field at the cortical level between 100 and 140 *V /m*. The location of the maximum electric field induced by TMS on the cortical surface was always kept on the convexity of the targeted gyrus with the induced current perpendicular to its main axis. Stimulation was delivered with an interstimulus interval jittering randomly between 2000 and 2300 ms (0.4–0.5Hz).

Stimulation coordinates were recorded and the electrodes positions were digitized. Trigeminal stimulation and muscle artefacts were minimized by placing the coil on a scalp area close to the midline, far away from facial or temporal muscles and nerve endings. To prevent contamination of TMS-evoked EEG potentials by the auditory response to the coil’s click, subjects wore earphones through which a noise masking, reproducing the time-varying frequency components of the TMS click, was played throughout each TMS/hd-EEG session. Our EEG amplifier (60 channels, 2 additional electrooculograms) uses a sample-and-hold mechanism to avoid the TMS induced artefact. In combination with the flat open ring carbon electrode design and the low impedance, it allows to recover a usable EEG signal 8 to 10 ms after the pulse.

In this study we did not perform a sham condition, as it was performed in previous studies using exactly the same setup as we used in our experiments, as well as in two other studies using a different setup. These studies showed that the TMS evoked potentials were absent in the sham condition, and that they were not confounded by auditory evoked potentials (Massimini et al., 2005; Rosanova et al., 2009, 2012; Ragazzoni et al., 2013).

Out of the initial 14 subjects, we excluded 5 of them for the precuneus and 2 for premotor, because of a low signal-to-noise ratio of TMS/EEG-evoked responses. TMS trials containing noise, muscle activity, or eye movements were detected and rejected (Rosanova et al., 2012). EEG data were average referenced, downsampled at half of the original sampling rate (from 725 Hz to 362 Hz), and bandpass filtered (2 to 80 Hz).

Source reconstruction was performed as in (Casali et al., 2013). Conductive head volume was modeled according to the 3-spheres BERG method (Berg and Scherg, 1994) as implemented in the Brainstorm software package (freely available at: http://neuroimage.usc.edu/brainstorm) and included three concentric spheres with different homogeneous conductivity, representing the best-fitting spheres of inner skull, outer skull and scalp compartments extracted from individual MRIs. The solution space was constrained to the cerebral cortex that was modeled as a three-dimensional grid of 3004 fixed dipoles oriented normally to cortical surface. This model was adapted to the anatomy of each subject using the Statistical Parametric Mapping software package (SPM8, freely available at: http://www.fil.ion.bpmf.ac.uk/spm) as follows: binary masks of skull and scalp obtained from individual MRIs were warped to the corresponding canonical meshes of the Montreal Neurological Institute (MNI) atlas. Then, the inverse transformation was applied to the MNI canonical mesh of the cortex for approximating to real anatomy.

Finally, EEG sensors and individual meshes were co-registered by rigid rotations and translations of digitized landmarks (nasion, left and right tragus). The single trial distribution of electrical sources in the brain was estimated by applying the empirical Bayesian approach as described in (Phillips et al., 2005; Mattout et al., 2006).

In order to summarize significant functional measures over anatomically and/or functionally identifiable brain regions, the time courses of the 3004 reconstructed sources were then averaged into the specific 90 cortical and subcortical areas of the Automated Anatomical Labeling (AAL) (Tzourio-Mazoyer et al., 2002) parcellation (Fig.1), according to their position on the cortical mesh.

**Figure 1:**
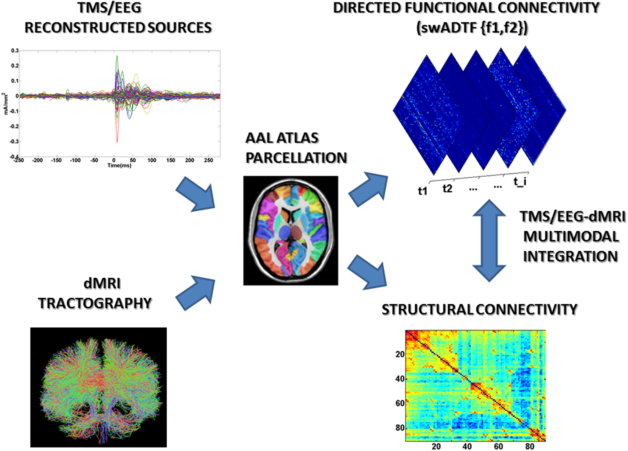
Flow chart of TMS/EEG-dMRI modeling. Up: the time courses of the 3004 reconstructed dipoles were averaged into the parcels of the Automated Anatomical Labeling (AAL) atlas (Tzourio-Mazoyer et al., 2002), consisting of 90 unique brain regions (cerebellar regions were excluded from the analysis). The 90 time courses obtained were modeled using spectrum-weighted adaptive directed transfer function (swADTF)(Van Mierlo et al., 2011, 2013). swADTF returns the causal interactions between the cortical regions (90*x*90 time varying directed functional connectivity matrices) at a specific frequency interval (f1,f2). Bottom: for each dMRI dataset whole-brain probabilistic tractography was performed using a combination of FSL and MRTRIX (see **Materials and methods**). The AAL atlas was then used to segment the fiber bundles between each pair of ROIs. Next, we determined the percentage of tracts between each pair of regions of the AAL template, resulting in a 90*x*90 structural connectivity matrix.

### dMRI data

#### Acquisition and preprocessing

A series of diffusion-weighted magnetic resonance images (dwi) of brain anatomy were acquired in each participant using a Siemens Trio Magnetom 3 Tesla system (Siemens Trio, University Hospital of Liege, Belgium). Diffusion-weighted images were acquired at a b-value of 1000 *s/mm*^2^ using 64 encoding gradients that were uniformly distributed in space by an electrostatic repulsion approach (Jones et al., 1999). Voxels had dimensions of 1.8 × 1.8 × 3.3 *mm*^3^ and volumes were acquired in 45 transverse slices using a 128 × 128 voxel matrix. A single T1-weighted 3D magnetization-prepared rapid gradient echo sequence (MPRAGE) image, with isotropic resolution of 1 *mm*^3^, was also acquired for each subject.

Diffusion volumes were analysed using typical preprocessing steps in dMRI (Zalesky et al., 2014; Caeyenberghs et al., 2012). Eddy current correction for each participant was achieved using FDT, v2.0, the diffusion toolkit within FSL 5.0 (FMRIBs Software Library; http://www.fmrib.ox.ac.uk/fsl). The eddy current correction step minimized distortions induced by eddy currents and also aligned each diffusion weighted volume to the first non-diffusion weighted volume to correct for simple intra-acquisition head movement. Rotations applied to the diffusion-weighted volumes were also applied to the corresponding gradient directions (Leemans and Jones, 2009). A fractional anisotropy (FA) image was estimated using weighted linear least squares fitted to the log-transformed data for each subject.

#### Registration of the anatomical image and atlas parcellation

We segmented each subject’s T1-weighted image into whole-brain white matter (WM), gray matter (GM), and cerebrospinal fluid (CSF) masks using FAST, part of FSL (FMRIB Software Library v 5.0). The corresponding white matter mask image was registered without resampling to the relevant dwi series (target image = thresholded FA image (FA *>* 0.2)) using FLIRT, v5.5, 12 degrees of freedom, nearest neighbour interpolation, mutual information cost function (Smith et al., 2004). The registration was performed without resampling in order to maintain the high spatial resolution of the structural image in the diffusion space.

As previously stated, the 90 cortical and subcortical nodes comprising the automated anatomical labeling (AAL) template (Tzourio-Mazoyer et al. 2002) were used as candidate atlas. The atlas was first registered to the T1 space using linear (FSL flirt) and non-linear warping (FSL FNIRT) in order to achieve the best registration into each subject’s space. Then, the single subject AAL template was finally registered without resampling to the dwi space using the affine transform resulting from the WM registration. This transformation matrix was also applied to the T1-derived GM mask which was used as termination mask for the tractography analysis.

#### Tractography and connectome construction

The fiber response model was estimated for each subject from the high b-value (b = 1000 s/mm2) diffusion-weighted images. A mask of single fiber voxels was extracted from the thresholded and eroded FA images. Only strongly anisotropic (FA *>* 0.7) voxels were used to estimate the spherical-harmonic coefficients of the response function (Tournier et al., 2004, 2008). Using non-negativity constrained spherical deconvolution, fiber orientation distribution (FOD) functions were obtained at each voxel using the MRTRIX3 package (J-D Tournier, Brain Research Institute, Melbourne, Australia, https://github.com/jdtournier/mrtrix3) (Tournier et al., 2012). For both the response estimation and spherical deconvolution steps we chose a maximum harmonic order *l*_*max*_ of 6.

Probabilistic tractography was performed using randomly placed seeds within subject-specific white matter masks, registered as mentioned in the latter. Fiber tracking settings were as follows: number of tracks = 10 million, FOD magnitude cutoff for terminating tracks = 0.1, minimum track length = 5 mm, maximum track length = 200 mm, minimum radius of curvature = 1 mm, tracking algorithm step size = 0.5 mm. Streamlines were terminated when they extended out of the WM-GM mask interface, or could not progress along a direction with an FOD magnitude or curvature radius higher than the minimum cutoffs.

The streamlines obtained were mapped to the relevant nodes defined by the AAL parcellation registered in the subject’s dwi space, using MRTRIX3 (Tournier et al., 2012). Each streamline termination was assigned to the nearest gray matter parcel within a 2 mm search radius. The resulting connectome was finally examined by determining the connection density (number of fiber connections per unit surface) between any two regions of the AAL template, as in (Caeyenberghs et al., 2012) (see also Fig.1). This correction was needed to account for the variable size of the cortical ROIs of the AAL template (Hagmann et al., 2008).

### TMS/hd-EEG directed functional connectivity estimation

#### Spectrum-weighted adaptive directed transfer function

Since we were interested in studying information transfer in the frequency domain, we evaluated directed functional connectivity using a multivariate model of spectral coefficients, i.e. the directed transfer function (DTF) (Kaminski and Blinowska, 1991; Kamiński et al., 2001; Babiloni et al., 2005). In order to cope with the non-stationary nature of the signals under study, we used the adaptive directed transfer function (ADTF) (Astolfi et al., 2008; Wilke et al., 2008). Specifically, we adopted the spectrum-weighted adaptive directed transfer function (swADTF)(Van Mierlo et al., 2013), which has been successfully used for connectivity modeling of epileptic intracranial EEG data (Van Mierlo et al., 2011, 2013).

A time-variant multivariate autoregressive (TVAR) model is built from the TMS/hd-EEG sources by using the Kalman filtering algorithm (Arnold et al., 1998; Schlögl et al., 2000; Van Mierlo et al., 2011). The time-variant connectivity measure, the swADTF, is calculated from the coefficients of the TVAR model as follows:

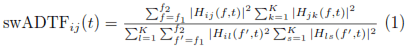

where *H*_*ij*_(*f, t*) in eq. 1 is the time-variant transfer matrix of the system describing the information flow from signal *j* to *i* at frequency *f* at time *t*, for each of the *K* signals. Each term *H*_*ij*_(*f, t*) is weighted by the autospectrum of the sending (in this case j) signal.

The swADTF allows us to investigate the causal relation between all the signals at a predefined frequency band over time. The measure weighs all outgoing information flow present in the terms *H*_*ij*_(*f, t*) by the power spectrum of the sending signal j. Each swADTF value corresponds to the directed time-variant strength of the information flow between two nodes. This dynamic interaction between nodes can also be represented as a series of time-varying directed matrices (see also Fig.1). The swADTF is normalized so that the sum of incoming information flow into a channel at each time point is equal to 1:

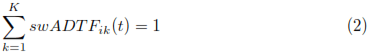

### TMS/EEG-dMRI multimodal integration

#### Outdegree computation and statistical assessment

We computed directed functional connectivity (swADTF) on the brain network defined by the anatomical atlas (AAL) reconstructed sources for each subject. Two parameters are needed for the swADTF calculation: the model order (*p*) and the update coefficient (*U C*). The model order defines how many previous time points are taken into account to update the dynamic interaction between nodes. The update coefficient defines how quickly the model will adapt to changes in the dataset. In this paper we set *p* = 5 and *UC* = 0.001. A detailed discussion on the implementation and the setup of the parameters can be found in (Van Mierlo et al., 2011).

The swADTF was calculated in 3 frequency bands: *α* (8-12 Hz), *β* (13-20 Hz), *β*2*/γ* (21-50 Hz). This choice followed the evidence that TMS on healthy awake subjects consistently evoked dominant EEG oscillations in different cortical areas (Rosanova et al., 2009). In particular, precuneus was shown to respond to TMS in the *β* band, the premotor area in the *β*2*/* γ and the occipital in *α*. These findings suggest that different brain areas might be normally tuned to oscillate at a characteristic rate (i.e. natural frequency)(Rosanova et al., 2009).

In order to track modulations of information flow due to TMS, we considered 2 different non-overlapping windows of 300 ms: a “baseline”, pre TMS stimulus, extended from 500 ms to 200 ms before the TMS pulse; a “post stimulus”, directly after TMS, which captures the dynamics from 20 to 300 ms after the pulse (the first 20 ms were discarded to minimize the effect of possible artifacts occurring at the time of stimulation, (Rogasch et al., 2013; Rosanova et al., 2009)).

We obtained the mean global outgoing flow from a region *j* before and after the stimulation by averaging the swADTF time courses in each of the two time windows and by summing the average amount of information transferred from *j* to each node of the network. In network terms, this quantity is called Outdegree. In our case, for each frequency band and window (i.e. baseline or post stimulus):

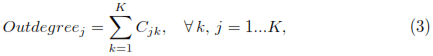

where *K* = 90 in our case (i.e. the number of AAL regions), and *C* is the connectivity matrix constructed by averaging the swADTF time courses within each window. All self-edges were set to 0. By using this procedure we aimed to obtain an illustrative snapshot of the total information flow from a region *j* at a specific stage of the TMS process (i.e. baseline or post stimulus).

In order to detect significant group changes in the Outdegree before and after the stimulation, a two-sample t-test of the post stimulus Outdegree against the correspondent baseline Outdegree was performed in each region. The choice of this statistical test over others was dictated by the fact that swADTF time courses are computed using a Kalman filtering algorithm, which assumes that error terms and noise are normally distributed (Van Mierlo et al., 2011; Arnold et al., 1998). Post stimulus Outdegree values were considered significant at *p* ≤ 0.05, False Discovery Rate (FDR) corrected for multiple comparisons (i.e. for *K* = 90 independent tests).

For each subject, the structural Outdegree of a node *j* (OutSC) was simply calculated from the structural connectivity matrix *S* by summing over its columns.

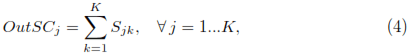

#### Structure-function correlations and statistical assessment

The dynamic interaction between regions modeled by swADTF can be represented as a series of time-varying directed connectivity matrices (see also Fig.1). In each frequency band, dynamic spatial correlation was defined as the mean rowby-row Pearson’s correlation at each time point between each subject’s directed functional connectivity matrix and the correspondent structural connectivity matrix.

The confidence intervals for the Pearson’s correlation distribution at the baseline were calculated by using a non-parametric bootstrap procedure (Efron and Tibshirani, 1986). The correlation coefficient was recomputed *n* = 100 times on the resampled data obtained by *n* random permutations of the values in the directed functional connectivity matrices at each time point of the baseline, while leaving the structural connectivity matrix unchanged. The empirical distribution of the resampled dynamic spatial correlation values at the baseline was used to approximate the sampling distribution of the statistic. A 95% confidence interval for the baseline was then defined as the interval spanning from the 2.5*th* to the 97.5*th* percentile of the obtained distribution. Values in dynamic spatial correlation that were falling outside this interval were considered significantly different from the baseline correlation.

To test for local structure-function interactions, we computed group-wise Spearman’s correlation between the cortical regions where the post stimulus Outdegree was significantly different from baseline and their correspondent OutSC value. Correlation was considered significant at *p* ≤ 0.05, where Spearman’s p-values were calculated using the exact permutation distributions for small sample sizes (Best and Roberts, 1975).

Following the hypothesis that TMS pulse gives rise to different connected cortical regions in the brain at different natural frequencies depending on the stimulated area (Rosanova et al., 2009), we evaluated structure-function correlation between flow information at the natural frequency and structural connectivity in three regions-of-interest (ROIs), previously defined and validated in (Rosanova et al., 2009) (i.e. occipital, precuneal and premotor area; Broadmann area 19, 7 and 6 respectively, see also Table 3 for a detailed list of the AAL regions included).

Dynamic spatial correlation was here evaluated by concatenating the swADTF time courses of each AAL region in the ROIs at its own natural frequency band (i.e. *α* for the occipital area, *β* for precuneus, *β*2*/γ* for premotor, as specified in (Rosanova et al., 2009)) and the corresponding fiber densities from the structural connectivity matrix. The confidence intervals were again calculated from the empirical baseline distribution using a non-parametric bootstrap procedure (Efron and Tibshirani, 1986), as formerly explained.

## Results

The dynamic spatial correlation between the directed functional connectivity (swADTF) and the connectome, for the two different sites of stimulation (i.e. left precuneus and left premotor) and for each of the three chosen frequency bands (i.e. *α*, *β*, *β*2*/γ*) deviates from baseline directly after the TMS pulse (Fig.2). This global network behavior does not depend on the subject or the stimulation site. The stable baseline configuration is then recovered after 200-300 ms, depending on the frequency band. Specifically, this temporary modification generated by TMS over the brain network is more pronounced (i.e. higher deviation from the baseline correlation) and faster in the *β*2*/γ* and *β* bands, while the return to baseline seems slower and less peaked in the *α* band.

**Figure 2:**
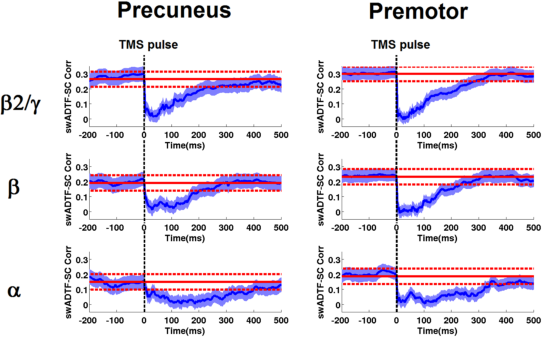
Time-varying spatial correlation between directed functional connectivity and structural connectivity. Each plot shows the average over subjects of the dynamic spatial correlation between the directed functional connectivity (swADTF) matrices and the structural connectivity (SC) in function of time (blue line, standard error in shaded blue), for the three different frequency bands (*α*, *β*, *β*2*/*γ, (Rosanova et al., 2009)). The red line indicates the mean baseline value, the dashed lines represent 95% confidence interval of the empirical baseline distribution (see **Materials and Methods**). Note the TMS-induced decrease in the observed structure-function correlation, for both stimulation sites and in each frequency band.

The significant differences (Table 1) in information flow across cortical regions after TMS perturbation are illustrated by projecting the OutDegree onto the anatomical template (Fig.3, see also the movies in **Supplementary Material**). The two sites of stimulation have peaks in information transfer at different frequency bands. In particular, the precuneus area has a maximum of information flow in the *β* band in proximity of the stimulation site, whereas the premotor has a maxima in the *β*2*/γ* band, more spread towards the hemisphere controlateral to the stimulation site.

**Table 1:** Percentage increase before/after stimulation for the AAL areas illustrated in Fig. 3 where information flow after TMS was significantly higher than baseline (*p <* 0.05, FDR corrected, see **Materials and Methods**), for the two sites of stimulation (left precuneus and left premotor) and the three different frequency band (F.B., i.e. *α*, *β*, *β*2*/γ*). Top increment for each site of stimulation is highlighted in bold.

| TMS SITE: LEFT PRECUNEUS |  |  | TMS SITE: LEFT PREMOTOR |  |  |
| --- | --- | --- | --- | --- | --- |
| AAL REGION | INCREASE F.B.<br>(%) |  | AAL REGION | INCREASE F.B.<br>(%) |  |
| Precentral<sub>L</sub> | 120 | $\alpha$ | Supp_Motor_Area<sub>L</sub> | 210 | $\alpha$ |
| Frontal_Mid<sub>L</sub> | 133 | $\alpha$ | Supp_Motor_Area<sub>R</sub> | 190 | $\alpha$ |
| Frontal_Inf_Oper<sub>L</sub> | 70 | $\alpha$ | Cingulum_Post<sub>R</sub> | 60 | $\alpha$ |
| Rolandic_Oper<sub>L</sub> | 60 | $\alpha$ | Cuneus<sub>L</sub> | 120 | $\alpha$ |
| Supp_Motor_Area<sub>L</sub> | 40 | $\alpha$ | Occipital_Sup<sub>L</sub> | 90 | $\alpha$ |
| Cingulum_Post<sub>R</sub> | 53 | $\alpha$ | SupraMarginal<sub>R</sub> | 50 | $\alpha$ |
| | | | Precuneus<sub>L</sub> | 80 | $\alpha$ |
| | | | Paracentral_Lobule<sub>R</sub> | 110 | $\alpha$ |
| Precentral<sub>L</sub> | 150 | $\beta$ | Frontal_Sup<sub>L</sub> | 185 | $\beta$ |
| Frontal_Mid<sub>R</sub> | 70 | $\beta$ | Frontal_Mid<sub>L</sub> | 150 | $\beta$ |
| Rolandic_Oper<sub>L</sub> | 50 | $\beta$ | Supp_Motor_Area<sub>L</sub> | 220 | $\beta$ |
| Supp_Motor_Area<sub>L</sub> | 90 | $\beta$ | Supp_Motor_Area<sub>R</sub> | 260 | $\beta$ |
| Parietal_Inf<sub>L</sub> | 167 | $\beta$ | Insula<sub>L</sub> | 50 | $\beta$ |
| **Precuneus<sub>L</sub>** | **280** | **$\beta$** | Cingulum_Mid<sub>R</sub> | 52 | $\beta$ |
| | | | Cingulum_Post<sub>R</sub> | 35 | $\beta$ |
| | | | Cuneus<sub>R</sub> | 30 | $\beta$ |
| | | | Cingulum_Mid<sub>R</sub> | 45 | $\beta$ |
| Precentral<sub>L</sub> | 233 | $\beta 2/\gamma$ | Precentral<sub>L</sub> | 118 | $\beta 2/\gamma$ |
| Frontal_Sup<sub>L</sub> | 152 | $\beta 2/\gamma$ | Frontal_Sup<sub>L</sub> | 363 | $\beta 2/\gamma$ |
| Supp_Motor_Area<sub>L</sub> | 126 | $\beta 2/\gamma$ | Frontal_Mid<sub>L</sub> | 203 | $\beta 2/\gamma$ |
| Supp_Motor_Area<sub>R</sub> | 130 | $\beta 2/\gamma$ | Frontal_Mid_Orb<sub>L</sub> | 153 | $\beta 2/\gamma$ |
| Cingulum_Mid<sub>L</sub> | 100 | $\beta 2/\gamma$ | Supp_Motor_Area<sub>L</sub> | 330 | $\beta 2/\gamma$ |
| Parietal_Inf<sub>L</sub> | 110 | $\beta 2/\gamma$ | **Supp_Motor_Area<sub>R</sub>** | **432** | **$\beta 2/\gamma$** |
| Precuneus<sub>L</sub> | 150 | $\beta 2/\gamma$ | Frontal_Sup_Medial<sub>R</sub> | 65 | $\beta 2/\gamma$ |
| | | | Insula<sub>L</sub> | 33 | $\beta 2/\gamma$ |
| | | | Cingulum_Mid<sub>R</sub> | 110 | $\beta 2/\gamma$ |
| | | | Thalamus<sub>L</sub> | 33 | $\beta 2/\gamma$ |
| | | | Temporal_Mid<sub>L</sub> | 53 | $\beta 2/\gamma$ |

**Figure 3:**
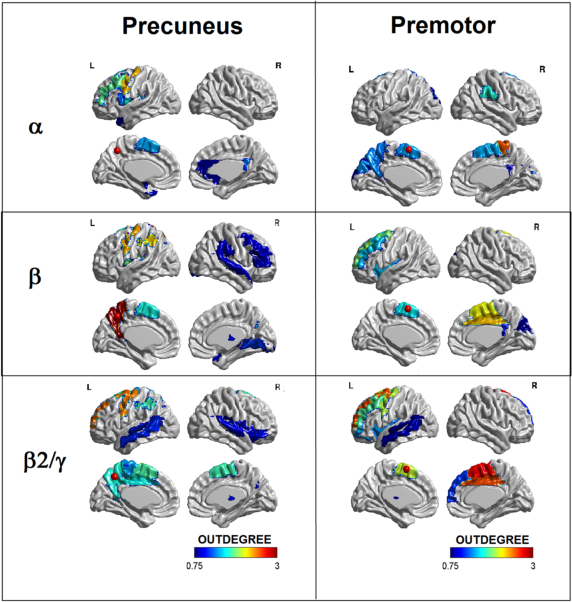
Information flow across cortical regions after TMS. Snapshot of differences between baseline and post TMS stimulus information transfer (i.e. Outdegree) at *p <* 0.05, FDR corrected (see **Materials and Methods**) across cortical regions, for the three predefined frequency bands (*α*, *β*, *β*2*/γ* (Rosanova et al., 2009)), obtained by averaging the swADTF time courses from 20 to 300 ms after the pulse. The red circles represent the stimulation site. Note that the precuneus area has a maximum of information flow in the *β* band in proximity of the stimulation site, whereas the premotor has a maxima in the *β*2*/γ* band, more spread towards the hemisphere controlateral to the stimulation site. These brain images were obtained using BrainNet Viewer (Xia et al., 2013).

Inter-subject Spearman’s correlation between the significant Outdegree values and their correspondent OutSC values also shows significant local peaks (Table 2) in proximity of the stimulation site (Fig.4), in the same frequency bands (i.e. *β* for precuneus and *β*2*/γ* for premotor). These results are in line with a previous study (Rosanova et al., 2009), where the authors showed that TMS on healthy awake subjects consistently evokes dominant oscillation in several cortical areas, particularly in 3 specific ROIs (i.e. occipital, precuneal and premotor area; Broadmann area 19, 7 and 6 respectively, see also Table 3). The evidence that each different brain area can be normally tuned by TMS to oscillate at a characteristic rate (i.e. natural frequency) might also explain the drop in structure-function correlation depicted in Fig.2. In fact, assuming that each of the 90 AAL cortical regions respond to TMS by oscillating at its peculiar natural frequency, the emergence of this complex between-band interaction might generate a consequent deflection in the within-band structure-function correlation (Fig.2). To further investigate this hypothesis, the dynamic spatial correlation between the directed functional connectivity (swADTF) of the 3 ROIs previously validated in (Rosanova et al., 2009) (see also Table 3) and the connectome, for both sites of stimulation, was calculated (Fig.5). As opposed to Fig.2, here for each ROI we evaluated the swADTF time course at its own natural frequency band (i.e. *α* for the occipital area, *β* for precuneus, *β*2*/γ* for premotor). The structure-function correlation significantly increases from the baseline, for both sites of stimulation (Fig.5). Notably, the increase is significant when the natural frequency bands of each ROIs are taken into consideration. On the other hand, this effect is not reproduced when considering frequency bands other than the natural ones for each ROI (Fig. S1 in **Supplementary Material**).

**Table 2:** List of local correlations (see **Materials and Methods**) after TMS for the AAL areas illustrated in Fig. 4, for the two sites of stimulation (left precuneus and left premotor) and the three different frequency band (F.B., i.e. *α*, *β*, *β*2*/γ*). The highest correlation values for each site of stimulation are highlighted in bold.

| TMS SITE: LEFT PRECUNEUS |  |  | TMS SITE: LEFT PREMOTOR |  |  |
| --- | --- | --- | --- | --- | --- |
| AAL REGION | $\rho$ | F.B. | AAL REGION | $\rho$ | F.B. |
| Precentral<sub>L</sub> | -0.60 | $\alpha$ | Supp_Motor_Area<sub>L</sub> | +0.40 | $\alpha$ |
| Frontal_Mid<sub>L</sub> | +0.41 | $\alpha$ | Supp_Motor_Area<sub>R</sub> | +0.31 | $\alpha$ |
| Frontal_Inf_Oper<sub>L</sub> | -0.34 | $\alpha$ | Cingulum_Post<sub>R</sub> | +0.33 | $\alpha$ |
| Rolandic_Oper<sub>L</sub> | +0.54 | $\alpha$ | Cuneus<sub>L</sub> | -0.60 | $\alpha$ |
| Supp_Motor_Area<sub>L</sub> | +0.32 | $\alpha$ | Occipital_Sup<sub>L</sub> | -0.32 | $\alpha$ |
| Olfactory<sub>R</sub> | +0.60 | $\alpha$ | Precuneus<sub>L</sub> | -0.55 | $\alpha$ |
| Cingulum_Ant<sub>R</sub> | -0.44 | $\alpha$ | Paracentral_Lobule<sub>R</sub> | +0.40 | $\alpha$ |
| Cingulum_Post<sub>R</sub> | +0.51 | $\alpha$ | Thalamus<sub>L</sub> | -0.53 | $\alpha$ |
| Temporal_Pole_Mid<sub>L</sub> | -0.36 | $\alpha$ | | | |
| Precentral<sub>L</sub> | +0.44 | $\beta$ | Precentral<sub>L</sub> | -0.41 | $\beta$ |
| Frontal_Inf_Orb<sub>R</sub> | +0.36 | $\beta$ | Frontal_Sup<sub>L</sub> | -0.34 | $\beta$ |
| Rolandic_Oper<sub>R</sub> | -0.52 | $\beta$ | Frontal_Mid<sub>L</sub> | -0.39 | $\beta$ |
| Supp_Motor_Area<sub>L</sub> | +0.35 | $\beta$ | Supp_Motor_Area<sub>L</sub> | -0.38 | $\beta$ |
| Supp_Motor_Area<sub>R</sub> | +0.38 | $\beta$ | Supp_Motor_Area<sub>R</sub> | +0.41 | $\beta$ |
| Insula<sub>R</sub> | +0.45 | $\beta$ | Insula<sub>L</sub> | +0.40 | $\beta$ |
| Cingulum_Mid<sub>L</sub> | -0.49 | $\beta$ | Cuneus<sub>R</sub> | +0.60 | $\beta$ |
| Parietal_Inf<sub>L</sub> | +0.35 | $\beta$ | Occipital_Sup<sub>R</sub> | +0.41 | $\beta$ |
| **Precuneus<sub>L</sub>** | **+0.65** | **$\beta$** | Occipital_Mid<sub>R</sub> | +0.50 | $\beta$ |
| Paracentral_Lobule<sub>L</sub> | +0.41 | $\beta$ | Occipital_Inf<sub>L</sub> | +0.54 | $\beta$ |
| Temporal_Sup<sub>R</sub> | -0.41 | $\beta$ | Occipital_Inf<sub>R</sub> | -0.56 | $\beta$ |
| Temporal_Mid<sub>L</sub> | +0.33 | $\beta$ | Temporal_Sup<sub>L</sub> | +0.37 | $\beta$ |
| | | | Temporal_Mid<sub>L</sub> | -0.32 | $\beta$ |
| | | | Temporal_Inf<sub>L</sub> | +0.54 | $\beta$ |
| Precentral<sub>L</sub> | -0.60 | $\beta 2/\gamma$ | Frontal_Sup<sub>L</sub> | +0.35 | $\beta 2/\gamma$ |
| Supp_Motor_Area<sub>L</sub> | -0.60 | $\beta 2/\gamma$ | Supp_Motor_Area<sub>L</sub> | +0.45 | $\beta 2/\gamma$ |
| Lingual<sub>R</sub> | +0.56 | $\beta 2/\gamma$ | **Supp_Motor_Area<sub>R</sub>** | **+0.64** | **$\beta 2/\gamma$** |
| Parietal_Inf<sub>L</sub> | +0.40 | $\beta 2/\gamma$ | ParaHippocampal<sub>L</sub> | +0.37 | $\beta 2/\gamma$ |
| Thalamus<sub>R</sub> | +0.51 | $\beta 2/\gamma$ | Calcarine<sub>R</sub> | +0.49 | $\beta 2/\gamma$ |
| Temporal_Sup<sub>R</sub> | -0.49 | $\beta 2/\gamma$ | Cuneus<sub>R</sub> | -0.42 | $\beta 2/\gamma$ |
| | | | Occipital_Sup<sub>L</sub> | -0.30 | $\beta 2/\gamma$ |
| | | | Thalamus<sub>L</sub> | -0.40 | $\beta 2/\gamma$ |
| | | | Thalamus<sub>R</sub> | -0.34 | $\beta 2/\gamma$ |
| | | | Temporal_Mid<sub>R</sub> | +0.37 | $\beta 2/\gamma$ |

**Figure 4:**
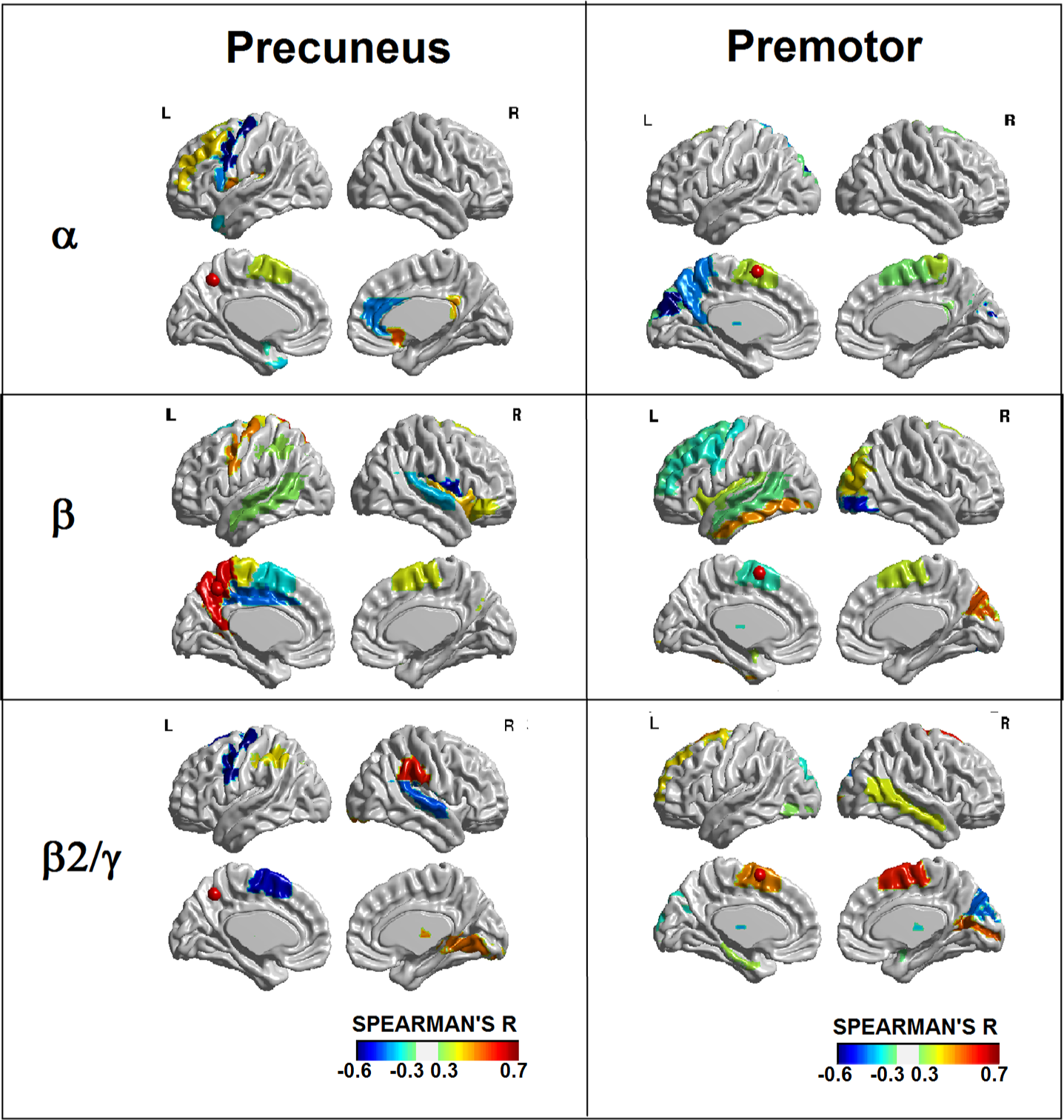
Local correlation between information flow and structural connectivity after TMS. Snapshot of the multi-subject Spearman’s correlation across cortical regions, for the three predefined frequency bands (*α*, *β*, *β*2*/γ* (Rosanova et al., 2009)), obtained by correlating regions with significant post stimulus Outdegree with their correspondent OutSC (see **Materials and Methods**). The red circles represent the stimulation site. These brain images were obtained using BrainNet Viewer (Xia et al., 2013).

**Table 3:** List of the AAL cortical areas included in the 3 regions-of-interest (i.e. occipital, precuneal and premotor area; Brodmann area (BA) 19, 7 and 6 respectively, see (Rosanova et al., 2009)) and corresponding natural frequency (N.F., (Rosanova et al., 2009)) selected for the analysis presented in Fig. 5.

| AAL REGION | ROI | BA | N.F. |
| --- | --- | --- | --- |
| Occipital_Sup_L | Occipital | 19 | $\alpha$ |
| Occipital_Sup_R | Occipital | 19 | $\alpha$ |
| Occipital_Mid_L | Occipital | 19 | $\alpha$ |
| Occipital_Mid_R | Occipital | 19 | $\alpha$ |
| Occipital_Inf_L | Occipital | 19 | $\alpha$ |
| Occipital_Inf_R | Occipital | 19 | $\alpha$ |
| Cingulum_Post_L | Precuneus | 7 | $\beta$ |
| Cingulum_Post_R | Precuneus | 7 | $\beta$ |
| Precuneus_L | Precuneus | 7 | $\beta$ |
| Precuneus_R | Precuneus | 7 | $\beta$ |
| Supp_Motor_Area_L | Premotor | 6 | $\beta 2/\gamma$ |
| Supp_Motor_Area_R | Premotor | 6 | $\beta 2/\gamma$ |
| Frontal_Mid_L | Premotor | 6 | $\beta 2/\gamma$ |
| Frontal_Mid_R | Premotor | 6 | $\beta 2/\gamma$ |

**Figure 5:**
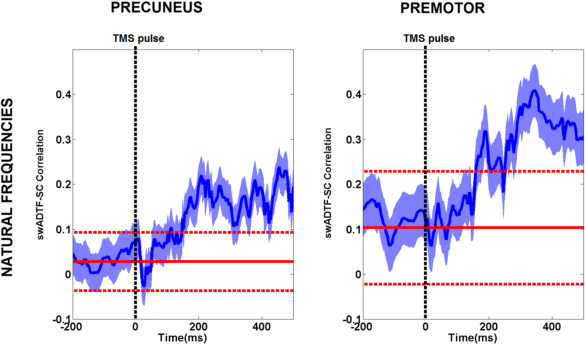
Time-varying spatial correlation at the natural frequency. Each plot shows the average over subjects of the dynamic spatial correlation (blue line, standard error in shaded blue) between the directed functional connectivity (swADTF) and structural connectivity (SC) for the ROIs defined in (Rosanova et al., 2009) (i.e. occipital, precuneus and premotor, see also Table 3), at their correspondent natural frequency bands (i.e. *α*, *β*, *β*2*/γ* respectively (Rosanova et al., 2009)), for both sites of stimulation. The continuous red line indicates the mean baseline value, the dashed lines represent 95% confidence interval of the empirical baseline distribution (see **Materials and Methods**). Note the increase in structure–function correlation for both sites of stimulation after TMS, when taking into account natural frequencies. This effect is not reproduced when considering frequency bands other than the natural ones for each ROI (Fig. S1 in **Supplementary Material**).

## Discussion

The relationship between structure and functional properties of brain networks at the macroscopic scale is a timely and challenging topic, currently investigated by means of different neuroimaging techniques (EEG, fMRI, dMRI) (Honey et al., 2010). In this work we studied the interplay between directed functional connectivity computed from TMS reconstructed EEG sources and the connectome extracted from whole-brain dMRI tractography. This is an unprecedented study of the relationship between brain structure (dMRI) and EEG dynamics using TMS as a probe of causal interaction.

It is known that stimulating peripheral receptors of different sensory systems results in evoked potentials with specific latencies, waveforms, and spectral components (Neidermeyer, 1999). TMS is also known to evoke electrical activations not only at the stimulated site but also in distant cortical regions (Ilmoniemi et al., 1997; Massimini et al., 2005). A previous study by Rosanova and colleagues revealed that distant areas, when activated by TMS, responded with oscillations closer to their own “natural” frequency (Rosanova et al., 2009). Nevertheless, the link between functional response to TMS and structural properties of the brain is still far from being clearly assessed.

As a first step, we compared structural and directed functional connectivity at the whole network level for different EEG bands (*α*, *β*, *β*2*/γ*). We observed a temporary decrease in the correlation between directed connectivity and structural connectivity after TMS. The extent and the duration of this deviation depended on the response of the brain network to the perturbation, involving different connected population of neural oscillators, each one with a characteristic operating frequency. In particular, we showed that, after stimulation, precuneus sends out information mostly in the *β* band, whereas premotor has peaks of information flow in the *β*2*/γ* band (Fig 3, Table 1). Assuming that the brain reacts to the perturbation with a complex pattern at mixed frequencies, then our findings suggest that the within-band deviation in the structure-function interplay after TMS might be caused by the rising between-band interactions in the whole brain network (Fig.2).

Our analysis on peaks of significant changes in information flow and local structure-function interactions at different frequency bands corroborated the hypothesis that TMS evokes dominant oscillation in different cortical areas at a characteristic rate (Rosanova et al., 2009). Each stimulated area appeared to mainly respond to the stimulation by sending the maximum amount information to the rest of the network in specific “natural” frequency bands, i.e. *β* for precuneus and *β*2*/γ* for premotor (Fig. 3, Table 1). Furthermore, the information sent from the stimulated region after the stimulation highly correlates with its structural connectivity (Fig. 4, Table 2), suggesting that the flow of information generated by cortical oscillations at different natural frequencies might be shaped and constrained by the structural architecture of the brain network. These empirical findings are in line with previous results in the field of theoretical brain network modeling (Marinazzo et al., 2012, 2014), where the authors showed that the maximum outgoing flow achievable for a brain region depends on its structural boundary.

These findings brought us to explore the link between the “natural” frequency response of some specific cortical areas and their structural architecture. We investigated the dynamic correlation between structure and function for the previously defined and validated ROIs in (Rosanova et al., 2009) (i.e. occipital, precuneal and premotor area; Broadmann area 19, 7 and 6 respectively, see also Table 3). Interestingly, the correlation between transfer of information at the natural frequency and structural connections increases after the stimulation and reveals a long-lasting effect over the selected ROIs (Fig.5). Moreover, this effect is not replicable when considering frequency bands other than the natural ones for each ROI (Fig. S1 in **Supplementary Material**). This might suggest that the interplay between cortical oscillators at specific resonant frequencies is directed and driven by their structural coupling (i.e. the amount of tracts connecting them).

These results lead to three main considerations. First, this work confirms the hypothesis that different rhythms in the brain emerge after TMS, and that this modulation is influenced by the structural connectivity among regions. This dynamic interaction at different natural frequencies seems to reflect intrinsic properties of cortical regions, and the way those are interconnected (Rosanova et al., 2009; Cona et al., 2011).

Secondly, our analysis permitted to evaluate the dynamic interactions between directed functional connectivity and anatomical connectivity, before and after TMS. The interplay between flow of information and structural connectivity at baseline is in line with findings reported in recent fMRI-dMRI studies (Barttfeld et al., 2015; Honey et al., 2009), where the rich repertoire of brain states do not necessarily correlate with the structural pattern. Here, our directed connectivity approach also allowed the investigation of the causal effects of systematic TMS-induced perturbations of the system, extending the insight on the relationship between structure and function. We showed that the information flow following the pulse is more connected to the structural pattern, when looking at natural frequencies (Fig. 5). This effect is not reproduced when considering frequency bands other than the natural ones for each ROI (Fig. S1 in **Supplementary Material**).

Thirdly, our multimodal whole-brain approach gives new insight on how TMS causally interferes with the brain network in healthy controls. More specifically, our study points out the importance of taking into account the major role played by different cortical oscillations when investigating the mechanisms for integration and segregation of information in the human brain (Balduzzi and Tononi, 2008; Casali et al., 2013). An interesting follow up of this study would indeed be to look at differences in structure-function interactions either when the cognitive function is pharmacologically modulated (i.e. anesthesia), or following pathology, damage or disruption in structural connections (i.e. coma and disorder of consciousness)(Fornito et al., 2015).

## Limitations

Given the intrinsic limitations of the EEG in terms of spatial resolution, it is important to stress that the patterns of connectivity detected by TMS/hd-EEG are necessarily coarse. Even though TEPs are characterized by a good testretest reproducibility (Lioumis et al., 2009), the inter–individual reproducibility of the outgoing flow of information could be improved by a better computation of the electric field induced by TMS. More advanced models (boundary, or finite, element models) could improve the accuracy of the source localization (Wagner et al., 2009).

We focused on TMS propagation through deep white matter pathways. It has been shown that TMS enhances the gray matter field around the site of stimulation (Opitz et al., 2011; Thielscher et al., 2011), and that certain superficial white matter systems pose challenges for measuring long-range cortical connections (Reveley et al., 2015). Future studies should also take into account the effect of the propagation electrical field through gray matter and superficial white matter fibers.

Another limitation of our study concerns the relatively small sample size and the inter-subject variability at the tractography level. In addition, it has been shown that there are many brain regions with complex fiber architecture, also referred to as crossing fibers (Jeurissen et al., 2011; Tournier et al., 2012). In this context, tractography approaches based on more advanced diffusion models (Jeurissen et al., 2011), or on more refined anatomical constraints (Smith et al., 2012) may provide more accurate anatomical connectivity patterns of brain networks. Therefore, our approach works best for studying large scale interactions than fine scale, local dynamics.

Finally, a b-value of 1000 *s/mm*^2^ is lower than the optimal one for performing CSD, about 2500-3000 *s/mm*^2^ (Tournier et al., 2013). However, despite of a low b-value, with a sufficient amount of directions crossing fibers can be reliably modeled with CSD and the result is still significantly better than with a simple DTI-based model, e.g. see (Roine et al., 2015) for a successful application.

## Conclusions

This work showed that different rhythms in the brain are evoked by TMS, and that this modulation is influenced by the structural connectivity among regions. We assessed that EEG directed functional connectivity induced by TMS is related to the underlying brain structure and to the frequency at which information is transferred. Crucially, in three specific cortical regions (precuneus, premotor, occipital), these frequencies coincide with the local predominant frequencies of TMS-induced activity. Our multimodal whole-brain analysis might offer new insights on how TMS causally interferes with the brain network in healthy controls, highlighting the importance of taking into account the dynamics of different local oscillations when investigating the mechanisms for integration and segregation of information in the human brain.

We thank Timo Roine, Erik Ziegler, Gianluca Frasso, Andrea Piarulli and Georgos Antonopoulos for the insightful discussion and comments on the manuscript. We thank Marie-Aurelie Bruno, Athena Demertzi, Audrey Vanhaudenhuyse and Melanie Boly for help in acquiring the data. This research was supported by the Wallonia-Brussels Federation of Concerted Research Action (ARC), Fonds National de la Recherche Scientifique de Belgique (FNRS), Belgian Science Policy (CEREBNET, BELSPO), McDonnell Foundation, European Space Agency, Mind Science Foundation, University Hospital and University of Liège. OB is a research fellow, OG a post doctoral fellow and SL a research director at FNRS.

## References

M. Arnold, X. Milner, H. Witte, R. Bauer, C. Braun, Adaptive ar modeling of nonstationary time series by means of kalman filtering. Biomedical Engineering, IEEE Transactions on 45(5), 553–562 (1998)

L. Astolfi, F. Cincotti, D. Mattia, F. de Vico Fallani, A. Tocci, A. Colosimo, S. Salinari, M.G. Marciani, W. Hesse, H. Witte, et al., Tracking the timevarying cortical connectivity patterns by adaptive multivariate estimators. Biomedical Engineering, IEEE Transactions on 55(3), 902–913 (2008)

F. Babiloni, F. Cincotti, C. Babiloni, F. Carducci, D. Mattia, L. Astolfi, A. Basilisco, P. Rossini, L. Ding, Y. Ni, et al., Estimation of the cortical functional connectivity with the multimodal integration of high-resolution eeg and fmri data by directed transfer function. Neuroimage 24(1), 118–131 (2005)

D. Balduzzi, G. Tononi, Integrated information in discrete dynamical systems: motivation and theoretical framework. PLoS computational biology 4(6), 1000091 (2008)

P. Barttfeld, L. Uhrig, J.D. Sitt, M. Sigman, B. Jarraya, S. Dehaene, Signature of consciousness in the dynamics of resting-state brain activity. Proceedings of the National Academy of Sciences 112(3), 887–892 (2015)

P.J. Basser, Inferring microstructural features and the physiological state of tissues from diffusion-weighted images. NMR in Biomedicine 8(7), 333–344 (1995)

P. Berg, M. Scherg, A multiple source approach to the correction of eye artifacts. Electroencephalography and clinical neurophysiology 90(3), 229–241 (1994)

D. Best, D. Roberts, Algorithm as 89: the upper tail probabilities of spearman’s rho. Applied Statistics, 377–379 (1975)

C. Bonato, C. Miniussi, P. Rossini, Transcranial magnetic stimulation and cortical evoked potentials: a tms/eeg co-registration study. Clinical neurophysiology 117(8), 1699–1707 (2006)

M. Bortoletto, D. Veniero, G. Thut, C. Miniussi, The contribution of tms–eeg coregistration in the exploration of the human cortical connectome. Neuroscience & Biobehavioral Reviews 49, 114–124 (2015)

E. Bullmore, O. Sporns, Complex brain networks: graph theoretical analysis of structural and functional systems. Nature Reviews Neuroscience 10(3), 186–198 (2009)

K. Caeyenberghs, A. Leemans, C. De Decker, M. Heitger, D. Drijkoningen, C.V. Linden, S. Sunaert, S. Swinnen, Brain connectivity and postural control in young traumatic brain injury patients: A diffusion mri based network analysis. Neuroimage: clinical 1(1), 106–115 (2012)

A.G. Casali, O. Gosseries, M. Rosanova, M. Boly, S. Sarasso, K.R. Casali, S. Casarotto, M.-A. Bruno, S. Laureys, G. Tononi, et al., A theoretically based index of consciousness independent of sensory processing and behavior. Science translational medicine 5(198), 198–105198105 (2013)

M. Catani, R.J. Howard, S. Pajevic, D.K. Jones, Virtual in vivo interactive dissection of white matter fasciculi in the human brain. Neuroimage 17(1), 77–94 (2002)

C. Chu, N. Tanaka, J. Diaz, B. Edlow, O. Wu, M. Hämäläinen, S. Stufflebeam, S. Cash, M. Kramer, Eeg functional connectivity is partially predicted by underlying white matter connectivity. Neuroimage (2014)

F. Cona, M. Zavaglia, M. Massimini, M. Rosanova, M. Ursino, A neural mass model of interconnected regions simulates rhythm propagation observed via tms-eeg. Neuroimage 57(3), 1045–1058 (2011)

N. De Geeter, G. Crevecoeur, A. Leemans, et al., Effective electric fields along realistic dti-based neural trajectories for modelling the stimulation mechanisms of tms. Physics in medicine and biology 60(2), 453 (2015)

B. Efron, R. Tibshirani, Bootstrap methods for standard errors, confidence intervals, and other measures of statistical accuracy. Statistical science, 54–75 (1986)

F. Ferrarelli, M. Massimini, S. Sarasso, A. Casali, B.A. Riedner, G. Angelini, G. Tononi, R.A. Pearce, Breakdown in cortical effective connectivity during midazolam-induced loss of consciousness. Proceedings of the National Academy of Sciences 107(6), 2681–2686 (2010)

A. Fornito, A. Zalesky, M. Breakspear, The connectomics of brain disorders. Nature Reviews Neuroscience 16(3), 159–172 (2015)

K.J. Friston, Another neural code? Neuroimage 5(3), 213–220 (1997)

M.S. George, Z. Nahas, M. Molloy, A.M. Speer, N.C. Oliver, X.-B. Li, G.W. Arana, S.C. Risch, J.C. Ballenger, A controlled trial of daily left prefrontal cortex tms for treating depression. Biological psychiatry 48(10), 962–970 (2000)

O. Gosseries, S. Sarasso, S. Casarotto, M. Boly, C. Schnakers, M. Napolitani, M.-A. Bruno, D. Ledoux, J.-F. Tshibanda, M. Massimini, et al., On the cerebral origin of eeg responses to tms: Insights from severe cortical lesions. Brain stimulation 8(1), 142–149 (2015)

S. Groppa, M. Muthuraman, B. Otto, G. Deuschl, H.R. Siebner, J. Raethjen, Subcortical substrates of tms induced modulation of the cortico-cortical connectivity. Brain stimulation 6(2), 138–146 (2013)

P. Hagmann, L. Cammoun, X. Gigandet, R. Meuli, C.J. Honey, V.J. Wedeen, O. Sporns, Mapping the structural core of human cerebral cortex. PLoS biology 6(7), 159 (2008)

S. Hofer, J. Frahm, Topography of the human corpus callosum revisitedcomprehensive fiber tractography using diffusion tensor magnetic resonance imaging. Neuroimage 32(3), 989–994 (2006)

C.J. Honey, J.-P. Thivierge, O. Sporns, Can structure predict function in the human brain? Neuroimage 52(3), 766–776 (2010)

C. Honey, O. Sporns, L. Cammoun, X. Gigandet, J.-P. Thiran, R. Meuli, P. Hagmann, Predicting human resting-state functional connectivity from structural connectivity. Proceedings of the National Academy of Sciences 106(6), 2035–2040 (2009)

R.J. Ilmoniemi, J. Virtanen, J. Ruohonen, J. Karhu, H.J. Aronen, T. Katila, et al., Neuronal responses to magnetic stimulation reveal cortical reactivity and connectivity. Neuroreport 8(16), 3537–3540 (1997)

B. Jeurissen, A. Leemans, D.K. Jones, J.-D. Tournier, J. Sijbers, Probabilistic fiber tracking using the residual bootstrap with constrained spherical deconvolution. Human brain mapping 32(3), 461–479 (2011)

D. Jones, M. Horsfield, A. Simmons, Optimal strategies for measuring diffusion in anisotropic systems by magnetic resonance imaging. Magn Reson Med 42 (1999)

M. Kamiński, M. Ding, W.A. Truccolo, S.L. Bressler, Evaluating causal relations in neural systems: Granger causality, directed transfer function and statistical assessment of significance. Biological cybernetics 85(2), 145–157 (2001)

M. Kaminski, K. Blinowska, A new method of the description of the information flow in the brain structures. Biological cybernetics 65(3), 203–210 (1991)

S. Komssi, S. Kähkönen, The novelty value of the combined use of electroencephalography and transcranial magnetic stimulation for neuroscience research. Brain research reviews 52(1), 183–192 (2006)

A. Leemans, D.K. Jones, The b-matrix must be rotated when correcting for subject motion in dti data. Magnetic Resonance in Medicine 61(6), 1336–1349 (2009)

P. Lioumis, D. Kičcić, P. Savolainen, J.P. Mäkelä, S. Kähkönen, Reproducibility of tmsevoked eeg responses. Human brain mapping 30(4), 1387–1396 (2009)

J.A. Maldjian, P.J. Laurienti, R.A. Kraft, J.H. Burdette, An automated method for neuroanatomic and cytoarchitectonic atlas-based interrogation of fmri data sets. Neuroimage 19(3), 1233–1239 (2003)

D. Marinazzo, G. Wu, M. Pellicoro, L. Angelini, S. Stramaglia, Information flow in networks and the law of diminishing marginal returns: evidence from modeling and human electroencephalographic recordings. PLoS one 7(9), 45026 (2012)

D. Marinazzo, M. Pellicoro, G. Wu, L. Angelini, J.M. Cortés, S. Stramaglia, Information transfer and criticality in the ising model on the human connectome. PLoS one 9(4), 93616 (2014)

M. Massimini, F. Ferrarelli, R. Huber, S.K. Esser, H. Singh, G. Tononi, Breakdown of cortical effective connectivity during sleep. Science 309(5744), 2228–2232 (2005)

J. Mattout, C. Phillips, W.D. Penny, M.D. Rugg, K.J. Friston, Meg source localization under multiple constraints: an extended bayesian framework. NeuroImage 30(3), 753–767 (2006)

E. Neidermeyer, The normal eeg of the waking adult. Electroencephalography: basic principles, clinical applications and related fields, 4th ed. Baltimore, MD: Williams and Wilkins, 149–173 (1999)

A. Opitz, M. Windhoff, R.M. Heidemann, R. Turner, A. Thielscher, How the brain tissue shapes the electric field induced by transcranial magnetic stimulation. Neuroimage 58(3), 849–859 (2011)

C. Phillips, J. Mattout, M.D. Rugg, P. Maquet, K.J. Friston, An empirical bayesian solution to the source reconstruction problem in eeg. NeuroImage 24(4), 997–1011 (2005)

A. Ragazzoni, C. Pirulli, D. Veniero, M. Feurra, M. Cincotta, F. Giovannelli, R. Chiaramonti, M. Lino, S. Rossi, C. Miniussi, Vegetative versus minimally conscious states: A study using tms-eeg, sensory and event-related potentials. PLoS one 8(2), 57069 (2013)

C. Reveley, A.K. Seth, C. Pierpaoli, A.C. Silva, D. Yu, R.C. Saunders, D.A. Leopold, Q.Y. Frank, Superficial white matter fiber systems impede detection of long-range cortical connections in diffusion mr tractography. Proceedings of the National Academy of Sciences, 201418198 (2015)

N.C. Rogasch, P.B. Fitzgerald, Assessing cortical network properties using tms–eeg. Human brain mapping 34(7), 1652–1669 (2013)

N.C. Rogasch, R.H. Thomson, Z.J. Daskalakis, P.B. Fitzgerald, Short-latency artifacts associated with concurrent tms–eeg. Brain stimulation 6(6), 868–876 (2013)

U. Roine, J. Salmi, T. Roine, T. Nieminen-von Wendt, S. Leppämäki, P. Rintahaka, P. Tani, A. Leemans, M. Sams, Constrained spherical deconvolution-based tractography and tract-based spatial statistics show abnormal microstructural organization in asperger syndrome. Molecular autism 6(1), 1–12 (2015)

M. Rosanova, A. Casali, V. Bellina, F. Resta, M. Mariotti, M. Massimini, Natural frequencies of human corticothalamic circuits. The Journal of Neuroscience 29(24), 7679–7685 (2009)

M. Rosanova, O. Gosseries, S. Casarotto, M. Boly, A.G. Casali, M.-A. Bruno, M. Mariotti, P. Boveroux, G. Tononi, S. Laureys, et al., Recovery of cortical effective connectivity and recovery of consciousness in vegetative patients. Brain, 340 (2012)

A. Schlögl, S. Roberts, G. Pfurtscheller, A criterion for adaptive autoregressive models, in Proceedings of the 22 nd IEEE International Conference on Engineering in Medicine and Biology, 2000, pp. 1581–1582

R.E. Smith, J.-D. Tournier, F. Calamante, A. Connelly, Anatomically constrained tractography: improved diffusion mri streamlines tractography through effective use of anatomical information. Neuroimage 62(3), 1924– 1938 (2012)

S.M. Smith, M. Jenkinson, M.W. Woolrich, C.F. Beckmann, T.E. Behrens, H. Johansen-Berg, P.R. Bannister, M. De Luca, I. Drobnjak, D.E. Flitney, et al., Advances in functional and structural mr image analysis and implementation as fsl. Neuroimage 23, 208–219 (2004)

S.-K. Song, S.-W. Sun, M.J. Ramsbottom, C. Chang, J. Russell, A.H. Cross, Dysmyelination revealed through mri as increased radial (but unchanged axial) diffusion of water. Neuroimage 17(3), 1429–1436 (2002)

O. Sporns, G. Tononi, R. Kötter, The human connectome: a structural description of the human brain. PLoS computational biology 1(4), 42 (2005)

A. Thielscher, A. Opitz, M. Windhoff, Impact of the gyral geometry on the electric field induced by transcranial magnetic stimulation. Neuroimage 54(1), 234–243 (2011)

J. Tournier, F. Calamante, D.G. Gadian, A. Connelly, et al., Direct estimation of the fiber orientation density function from diffusion-weighted mri data using spherical deconvolution. Neuroimage 23(3), 1176–1185 (2004)

J. Tournier, C.-H. Yeh, F. Calamante, K.-H. Cho, A. Connelly, C.-P. Lin, et al., Resolving crossing fibres using constrained spherical deconvolution: validation using diffusion-weighted imaging phantom data. NeuroImage 42(2), 617–625 (2008)

J. Tournier, F. Calamante, A. Connelly, et al., Mrtrix: diffusion tractography in crossing fiber regions. International Journal of Imaging Systems and Technology 22(1), 53–66 (2012)

J. Tournier, F. Calamante, A. Connelly, et al., Determination of the appropriate b value and number of gradient directions for high-angular-resolution diffusion-weighted imaging. NMR in Biomedicine 26(12), 1775–1786 (2013)

N. Tzourio-Mazoyer, B. Landeau, D. Papathanassiou, F. Crivello, O. Etard, N. Delcroix, B. Mazoyer, M. Joliot, Automated anatomical labeling of activations in spm using a macroscopic anatomical parcellation of the mni mri single-subject brain. Neuroimage 15(1), 273–289 (2002)

P. Van Mierlo, E. Carrette, H. Hallez, K. Vonck, D. Van Roost, P. Boon, S. Staelens, Accurate epileptogenic focus localization through time-variant functional connectivity analysis of intracranial electroencephalographic signals. Neuroimage 56(3), 1122–1133 (2011)

P. Van Mierlo, E. Carrette, H. Hallez, R. Raedt, A. Meurs, S. Vandenberghe, D. Roost, P. Boon, S. Staelens, K. Vonck, Ictal-onset localization through connectivity analysis of intracranial eeg signals in patients with refractory epilepsy. Epilepsia 54(8), 1409–1418 (2013)

A.N. Voineskos, L.J. O’Donnell, N.J. Lobaugh, D. Markant, S.H. Ameis, M. Niethammer, B.H. Mulsant, B.G. Pollock, J.L. Kennedy, C.F. Westin, et al., Quantitative examination of a novel clustering method using magnetic resonance diffusion tensor tractography. Neuroimage 45(2), 370–376 (2009)

A.N. Voineskos, F. Farzan, M.S. Barr, N.J. Lobaugh, B.H. Mulsant, R. Chen, P.B. Fitzgerald, Z.J. Daskalakis, The role of the corpus callosum in transcranial magnetic stimulation induced interhemispheric signal propagation. Biological psychiatry 68(9), 825–831 (2010)

T. Wagner, J. Rushmore, U. Eden, A. Valero-Cabre, Biophysical foundations underlying tms: setting the stage for an effective use of neurostimulation in the cognitive neurosciences. Cortex 45(9), 1025–1034 (2009)

C. Wilke, L. Ding, B. He, Estimation of time-varying connectivity patterns through the use of an adaptive directed transfer function. Biomedical Engineering, IEEE Transactions on 55(11), 2557–2564 (2008)

M. Xia, J. Wang, Y. He, et al., Brainnet viewer: a network visualization tool for human brain connectomics. PLoS one 8(7), 68910 (2013)

A. Zalesky, H. Akhlaghi, L.A. Corben, J.L. Bradshaw, M.B. Delatycki, E. Storey, N. Georgiou-Karistianis, G.F. Egan, Cerebello-cerebral connectivity deficits in friedreich ataxia. Brain Structure and Function 219(3), 969–981 (2014)

